# Difference in somatosensory event-related potentials in the blind subjects leads to better performance in tactile P300 BCI

**DOI:** 10.1101/2020.06.16.155796

**Authors:** Rafael Grigoryan, Dariya Goranskaya, Andrey Demchinsky, Ksenia Ryabova, Denis Kuleshov, Alexander Kaplan

## Abstract

In this study, we have created an 8-command P300 tactile BCI with two stimuli types, running on a minimally modified consumer Braille display and tested it on 10 blind subjects and 10 sighted controls. Blind subjects have demonstrated 27% higher median accuracy than sighted controls (*p* < 0.05), proving that the blind subjects are not only able to use tactile BCI but also can achieve superior results in comparison with sighted subjects. Median accuracy in the blind group with the best stimuli type has reached 95%. The difference in event-related potentials between groups is located in frontocentral sites before 300 ms post-stimulus and corresponds with early cognitive ERP components. The blind subjects have higher amplitude and lower latency of ERPs. This result is consistent through experimental conditions with different tactile stimuli. The classification performance for the blind subjects is correlated with Braille reading speed. This enables a discussion about mechanisms of plastic changes during sensory compensation after vision loss and its dependence on personal perceptual experience.

**Author summary:** Sensory compensation following vision loss can be recognized as a unique model for neural plasticity. However, the magnitude of the effect and the specific tasks where it’s manifested is still a subject for debate. In this study, we have created a tactile brain-computer interface game to study how somatosensory processing is different between the blind and the sighted people. The participants were required to attend to tactile stimuli, and the correct stimulus was selected using realtime EEG classification. We have shown, for the first time, that the blind subjects are significantly better than the sighted in tactile brain-computer interface tasks. We have also found, that individual performance is correlated with Braille proficiency. This result links personal perceptual abilities in two different sensory tasks. EEG analysis revealed that differences in performance can be attributed to early cognitive processing steps. Along with practical considerations in brain-computer interface development, the results also add to the data on cognitive processing in the blind and enable the discussion on the importance of Braille education.

## Introduction

Noninvasive brain-computer interfaces (BCIs) are routinely used for communication in disabled patients. Most widespread communicative BCIs employ paradigms based on various visual evoked potentials, including P300, SSVEP, and C-VEP. The subjects are supposed to direct their attention to the target of interest, which is activated in a manner to generate event-related potentials, that are classified to form a command to the computer. Visual P300 ERP paradigm is probably the most popular, being featured in various commercial products, enabling communication in patients, suffering from the inability to speak or move [1]. Most visual P300 BCI implementations rely on the direction of eye gaze and can become unusable for patients, who are not able to voluntarily control eye movement. However, the solutions for this problem are being proposed, with gaze-independent spellers based on rapid serial visual presentation [2, 3]. On the other hand, in the case of a patient with severely impaired or absent vision, it’s not possible to use most visual BCIs, creating a need for other sensory modalities, like auditory or tactile.

Tactile BCIs are not as well-studied as visual ones. This may be attributed to the necessity to create a dedicated stimulation device, which is more complicated than to use a computer screen. Another concern is lower performance characteristics that are typically achieved by tactile BCIs. This was demonstrated in [4], where participants were able to achieve near 100% target selection accuracy in visual P300 speller, while their accuracy in tactile P300 BCI never exceeded 80%. However, more recent studies [5] report subjects achieving accuracy over 95%, being able to control a wheelchair with a tactile array of powerful linear tactors, primarily used for haptic feedback vests. Other possible stimulation devices for tactile P300 BCI are electrical stimulators. There has been a report of P300 BCI, based on selective attention to one of the four fingers being electrically stimulated with the FES device [6]. The case of successfully decoding evoked potentials, produced by stimulation of a single finger with two different Braille-like patterns was also reported, with accuracy getting as high as 90% [7, 8]. However, with all this variety, refreshable Braille displays were not used as tactile stimulators in BCI.

Braille displays are readily available haptic devices that are already distributed to some extent within a low vision demographic. The creation of tactile BCI that can run on unaltered Braille displays can add to the distribution of the technology. Sensory properties of Braille cells are familiar to display users and their ergonomics have been studied extensively [9]. Cells can deliver various levels of force [10] and up to 256 different patterns. The studies, using Braille cells for stimulus presentation usually require creating a custom stimulator [4, 11].

The common feature of the abovementioned studies is the demography of the participants, being mostly healthy volunteers. This imbalance in BCI research has been discussed for a long time [12]. Most subjects in the research are either students or faculty staff, with no obvious health problems. Tests in the elderly people are rare [5], and tests in target demographics, like ALS patients, are even rarer [13]. This approach to subject inclusion may be efficient when the research is centered around the development of basic signal processing pipelines, however, when the resulting technology is rolled into production, unexpected problems may arise. For example, it is known that the ability of ALS patients to endure high cognitive workloads is inferior to those in healthy subjects [14], and the design of the spellers should be adjusted for successful use. The same issues are present in tactile BCIs. The primary justification for research in this area is the need to accommodate the needs of low vision patients (although there are attempts to use tactile BCI as a training tool for other patients [15]). However, to this day no tactile BCI was evaluated with the blind subjects.

Along with practical implications, a study like this may be useful for basic research, due to particular neurophysiological qualities of people, suffering from vision loss. BCI technology allows investigating complex psychophysiological issues in a gamified manner, thus reducing the effects of fatigue that are almost inevitable in lengthy neurophysiological experiments. Blindness is a convenient model for neural plasticity in humans, offering a unique opportunity to study how sensory experience influences the functional and anatomical properties of the brain. The results from neuroimaging studies with the blind subjects can be used to generate new insights into what localization of brain function exactly means. The anatomical areas that were routinely considered to be hard-wired may demonstrate consistent function only in typical subjects with typical sensory experience and can be significantly reorganized in subjects, suffering long-term vision loss [16]. The degree of this rearrangement, its dependence on various sensory experience and the connection between neuroanatomical structure and function provides a broad field of study. For example, it is important to access the functional differences in tactile perception that are present in the blind subjects. Although sensory compensation in the blind is a long-established phenomenon [17], the extent and particular characteristics of differences between the blind and the sighted cohorts are still subject to discussion. For example, the blind subjects are shown to perform better in the grating orientation task [18] and recently in a texture discrimination task [19]. However, some previous studies have claimed to find no difference between the blind and the sighted in tactile tests [20, 21], including the aforementioned [19], where groups did not differ in a shape discrimination task.

It is also still not obvious whether Braille reading practice influences performance. There were reports of the Braille readers being not better in tactile tests than the blind non-readers [18, 22]. Some reports even propose that Braille readers are worse in some tasks, for example [23], where the blind Braille readers were mislocating stimuli between hands and fingers more often than non-readers. This obviously can influence the viability of tactile BCI, relying on discrimination between fingers. On the other hand, practice is shown to increase tactile acuity, for example in musicians [24], and frequent Braille readers do have several hours of relevant tactile practice every day. It remains to be seen, whether Braille reading proficiency is correlated with tactile acuity and how this connection can be captured in different tasks.

## Materials and methods

### Subjects

11 peripherally blind and 10 sighted participants took part in the experiment. 5 blind subjects were congenitally blind, and 5 have been suffering vision loss for at least 8 years. 3 subjects had residual light sensitivity. Blind subjects were Braille-literate. One blind subject was excluded from analysis due to revealed medical condition, resulting in a high difference in tactile acuity between hands. 10 sighted subjects had normal or corrected-to-normal vision acuity and no known history of neurological disease. See Fig 1 for subject demographics. The study was performed in accordance with the declaration of Helsinki and all participants gave written informed consent.

**Fig 1.**
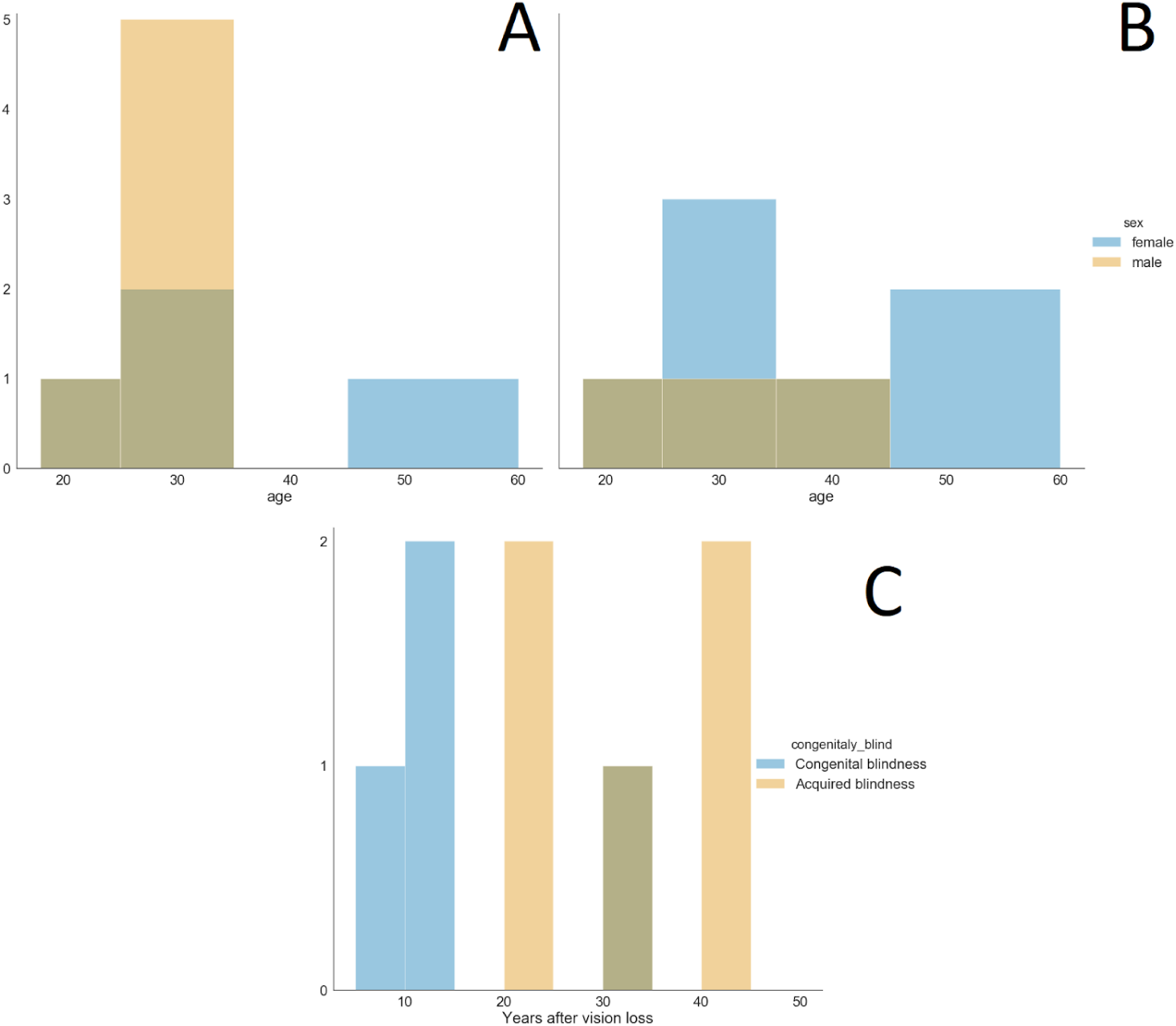
Subject demographics. (A) age and sex in the sighted group, (B) age and sex in the blind group, (C) time since vision loss in the blind group.

### Apparatus

EEG was recorded at 500 Hz using 45 Ag/AgCl gel electrodes and NVX-52 amplifier (MKS, Zelenograd, Russia). The electrodes used were: FP1, FP2, F3, Fz, F4, FC5, FC3, FC1, FCz, FC2, FC4, FC6, T7, C5, C3, C1, Cz, C2, C4, C6, T8, CP5, CP3, CP1, CPz, CP2, CP4, CP6, P7, P5, P3, P1, Pz, P2, P4, P6, P8, PO7, PO3, POz, PO4, PO8, O1, Oz, and O2. Electrodes were referenced to linked earlobes with AFz ground. Additionally, the ECG electrode was placed on the subject’s right wrist.

The stimulation device consisted of unaltered commercially available 40-cells Braille display ALVA USB 640 Comfort (Optelec, Barendrecht, the Netherlands). The device was connected via Bluetooth and controlled via BrlAPI [25]. Three rightmost Braille cells were utilized for custom optical synchronization device, generating TTL pulse when underlying cells switched into the activated position (Fig 2) Every stimuli activation that was presented to the subject was accompanied by synchronous activation of these cells and following the TTL pulse, which was detected by the EEG device and synced with the data stream. The EEG and TTL data was streamed to BCI application using LSL protocol [26]. The custom BCI application was written in Python 2.7, running on Windows 8 - based laptop.

**Fig 2.**
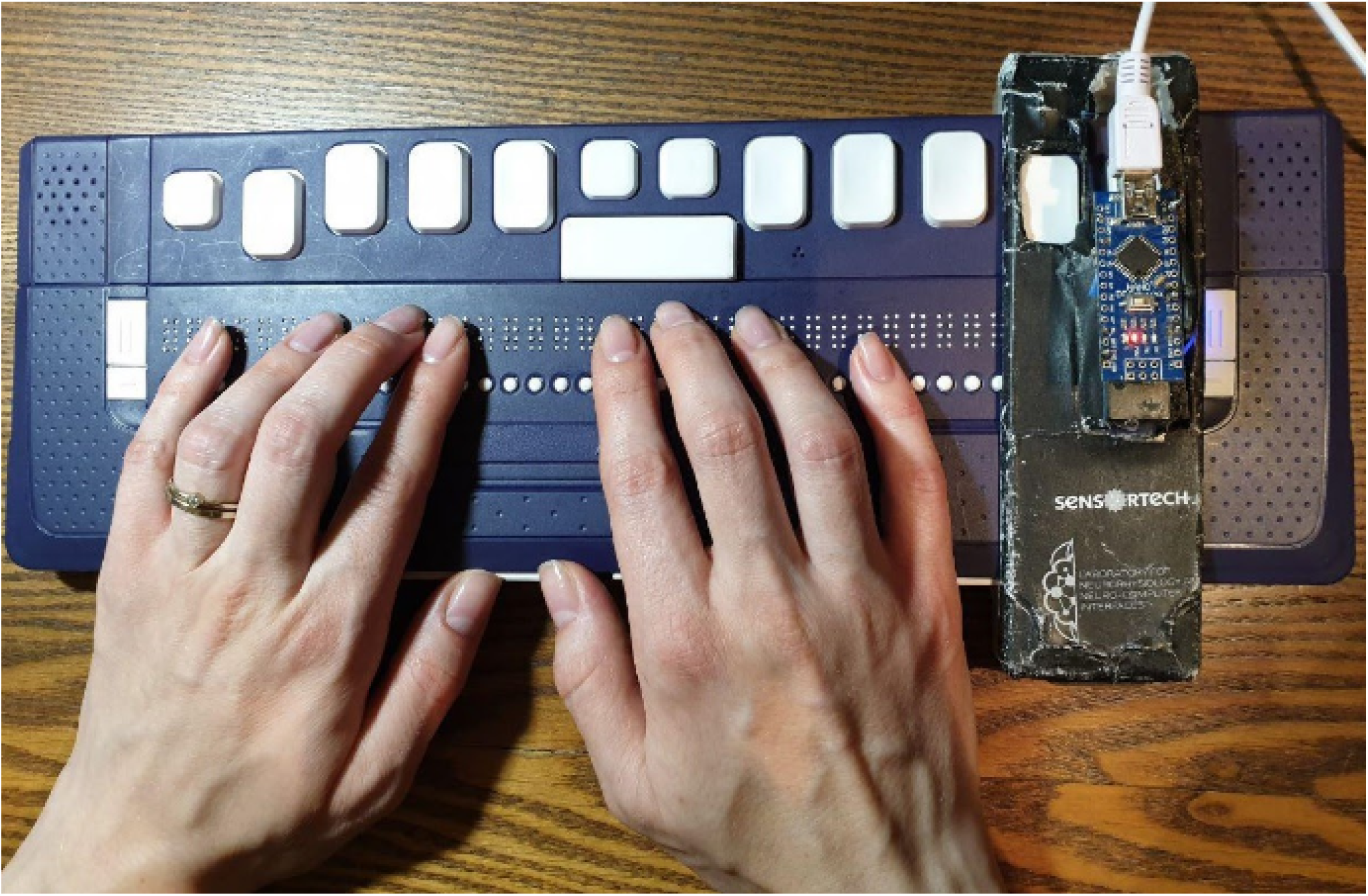
Tactile stimulation device. Subject’s hands are positioned on the Braille display. The synchronization device is on the right.

### Design and procedure

Before the experiment, subjects were required to read the text of the informed consent (around 5000 characters with spaces) in Russian. For the blind subjects, the text of informed consent was embossed using Braille printer (7 one-sided A4 pages), and the time spent on reading the text was used as a measure of Braille proficiency. Also, the subjects were asked to report the average time they spend reading Braille text (hours per week).

After installing the electrodes, the subjects were seated in front of the table and placed their hands on the Braille display. The task for the participant was to discriminate between the activation of Braille cells. All sessions used 8 active cells, so participants needed to use 8 fingers (index, ring, the middle, and pinky finger on each hand). To ensure comfort and accommodate different sized hands, the positions of active cells were adjusted for every participant. The stimulation consisted of activation of Braille cell (full length) for 200 ms with stimulus onset asynchrony of 300 ms. The Braille pattern was the same for all cells. The subject was instructed to count activations under the specific finger, and ignore all others. The target finger was provided to the participant using tactile cues before each session. The session consisted of 10 randomized activations of stimuli under all 8 fingers, after which the data was sent to the classifier. Each condition had 16 sessions, with every finger being the target for 2 times.

Two experimental conditions were using different stimulus size. The condition with large stimuli used activation of the full Braille cell (8 dots), and condition with small stimuli used activation of a single pin at the center of the cell (pin 5). Along with these two BCI sessions, participants optionally completed one or two other 10-15 minute long tasks with Braille display, which are not part of this study. All tasks were presented to the participant in a randomized order.

Each experimental condition began with the learning stage, which consisted of 6 sessions. This resulted in 60 target and 420 non-target trials (and corresponding feature vectors) being used to train the classifier. Then, the newly trained classifier was used to deliver feedback on selected targets via voice synthesizer for 10 next sessions. Participants were told that the aim of the game is to get the computer to guess the correct target.

For the sighted subjects, during the BCI sessions, the view of the Braille display was obstructed with a cardboard box.

### Data analysis

Recorded EEG was re-referenced using common average reference. Then, ICA was used to reject oculomotor artifacts. Since ICA is sensitive to low-frequency drifts, raw EEG was filtered using the 4th-order Butterworth filter from 1 to 35 Hz. Then, components, highly correlated with FP1 and FP2 channels, were rejected. The resulting unmixing matrix was applied to EEG, filtered from 0.1 to 35 Hz with an additional zero-phase notch filter at 50 Hz. This way, at the cost of slightly worse oculomotor artifact subtraction, we can preserve more accurate ERP waveforms, which are reported to suffer from excessive high-pass filtering [27, 28]. Then, filtered EEG was cut into epochs and averaged, with average evoked potentials being baseline-corrected using 50 ms pre-stimulus. Nontarget evoked potentials were subtracted from target ones to get rid of non-task-related activity. The resulting differential potential was used for analysis.

ERP statistics were carried out using the non-parametric spatiotemporal cluster-based permutation test with 10000 permutations, as described in [29, 30]. F-test was used as an estimator with an adaptive cluster threshold corresponding to a 0.05 significance level. The connectivity matrix was computed using Delaunay triangulation based on 2D sensor locations. The single average waveform for the cluster-based test was produced as the average differential potential for a single input trial. This method ensures multiple comparison correction by design, so no additional steps were taken. EEG processing and statistics used MNE-python 0.19 [31].

Both online classification during the experiment and offline accuracy estimation were based on the same classifier core. The preprocessing pipeline for offline classification was the same as for waveform analysis, with the difference in bandpass filter being 1 to 20 Hz. Narrower filter allowed for better signal-to-noise ratio. The adverse effects of excessive filtering are not so important during classification since the only difference we care about is the difference between target and non-target stimuli. Also, ICA components rejection was present only in offline mode. The data was cut into 800 ms epochs and downsampled by the factor of 10 with an anti-aliasing filter applied. Then, the subset of 11 channels was chosen (C3, Cz, C4, C6, CP3, CPz, CP4, Pz, PO3, PO4) and epochs were converted into feature vectors.

Feature vectors were classified using Fisher’s linear discriminant analysis with automatic Ledoit-Wolf shrinkage [32] from the Scikit-learn package [33]. The classifier returned the probability of each epoch being of the target class. Then the probabilities were multiplied across stimuli presentations, and the highest yielding stimuli were considered to be the target.

Offline accuracy was simulated with 1000-times stratified shuffle split cross-validation with classifier parameters similar to the online sessions. The scoring function did take into account dataset imbalance towards non-target responses, so the resulting numbers fully replicate experimental accuracy, with the exception of order effects in ERPs [34]. Pairwise comparisons of offline classification accuracies were carried out with the Mann-Whitney test. Prior to comparisons, the outliers, defined as 1.5 interquartile range, were removed from comparison. For within-subject comparisons between large and small stimuli, we were using the Wilcoxon signed-rank test. Correlation coefficients were computed using the Kendall tau statistic. All analysis steps were performed using Python 3.7.

## Results

### EEG

The evoked activity in the blind and the sighted subjects is dominated by a P300 peak around 300-400 ms. ERPs also have positivity before P300 in frontocentral sites, peaking around 150 ms post-stimulus in the blind subjects and around 300 ms in the sighted ones (see Fig 3 and Fig 4). This early positive component exhibits significant differences between the blind and the sighted both in condition with large and small stimuli.

**Fig 3.**
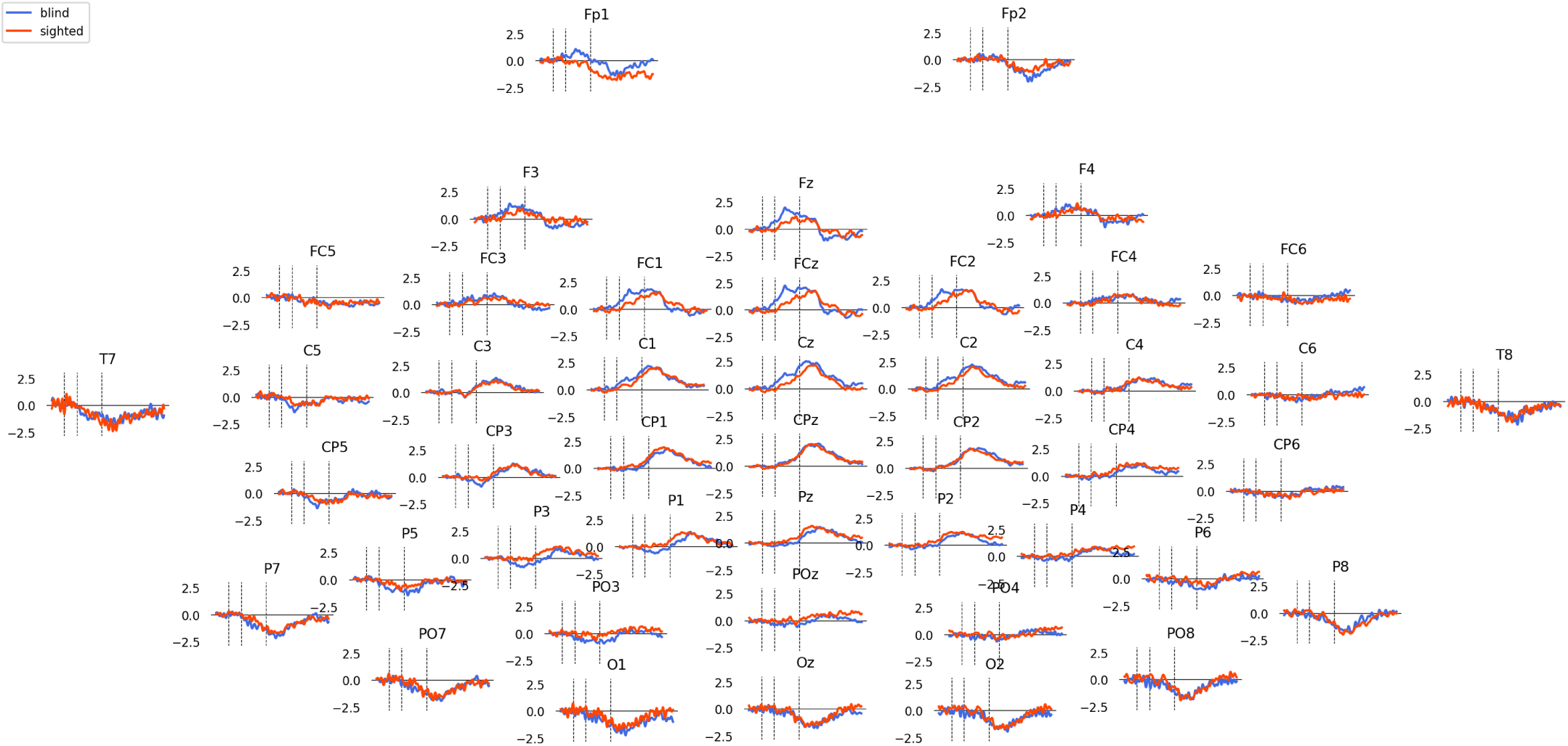
Grand average differential waveforms for the blind and the sighted subjects in condition with large stimuli. Vertical dashed lines represent 0, 100 and 300 ms post-stimulus.

**Fig 4.**
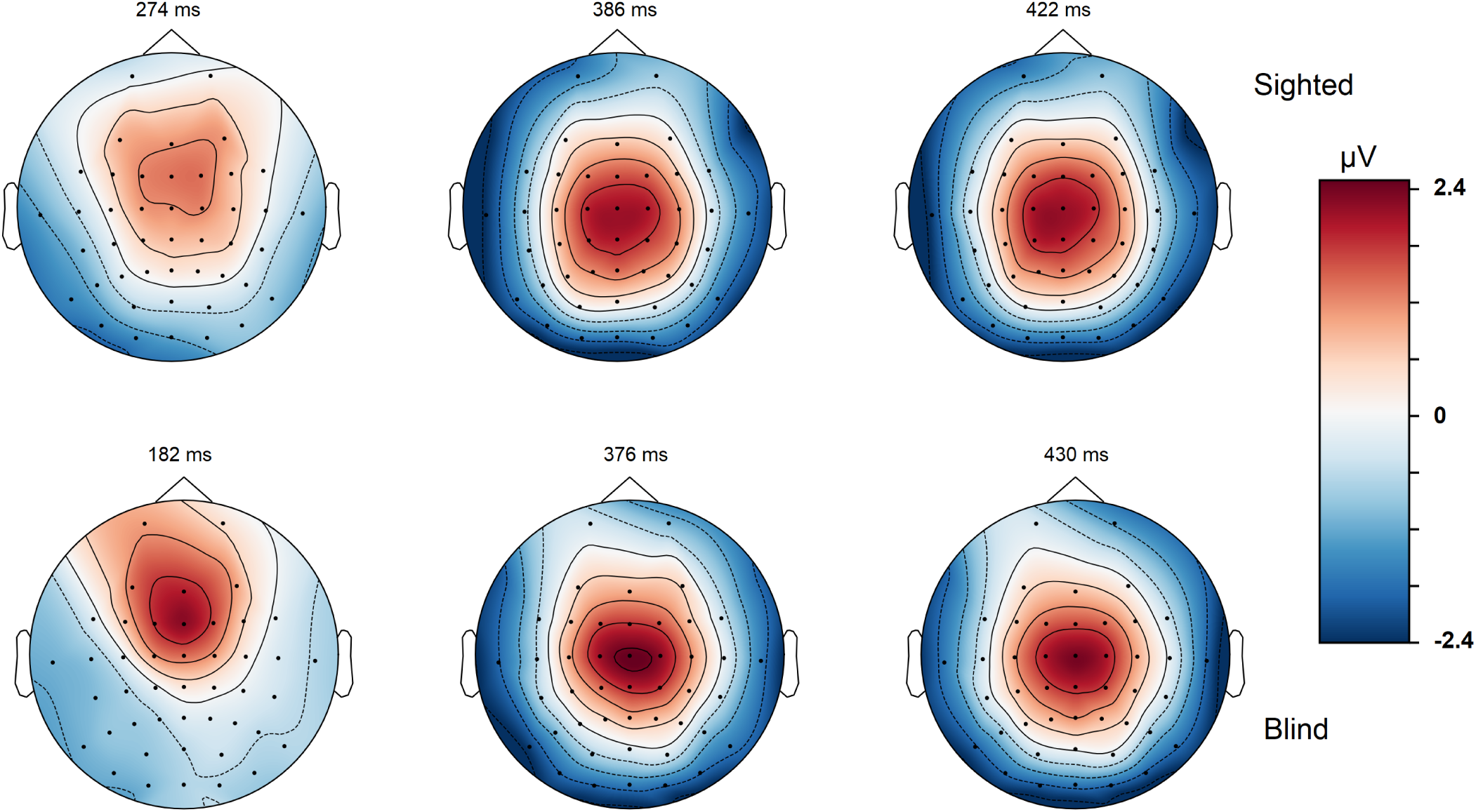
Topographical maps of evoked data for the blind and the sighted subjects. Peaks are selected automatically.

For large stimuli, the mean F-score for the significant cluster is 3.3 and *p* = 0.011. The cluster is located at central F, FC, and C electrode sites and the area of significance spans for approximately 200 ms starting around 60 ms poststimulus. The condition with smaller stimuli demonstrates a similar effect: significant frontocentral cluster from around 120 to 300 ms with a mean *F* = 3.55 and *p* < 0.03. See Fig 5 for details. Note that the existence of a significant cluster as a whole does not imply a significant difference at any specific time and site inside that cluster [30]. No significant differences were found for differential ERPs from right and left hands, or different fingers (including primary reading finger vs others). Also, no differences were found between the congenitally blind and the rest of the blind. Nontarget ERPs from the right and left hand demonstrate expected contralateral distribution.

**Fig 5.**
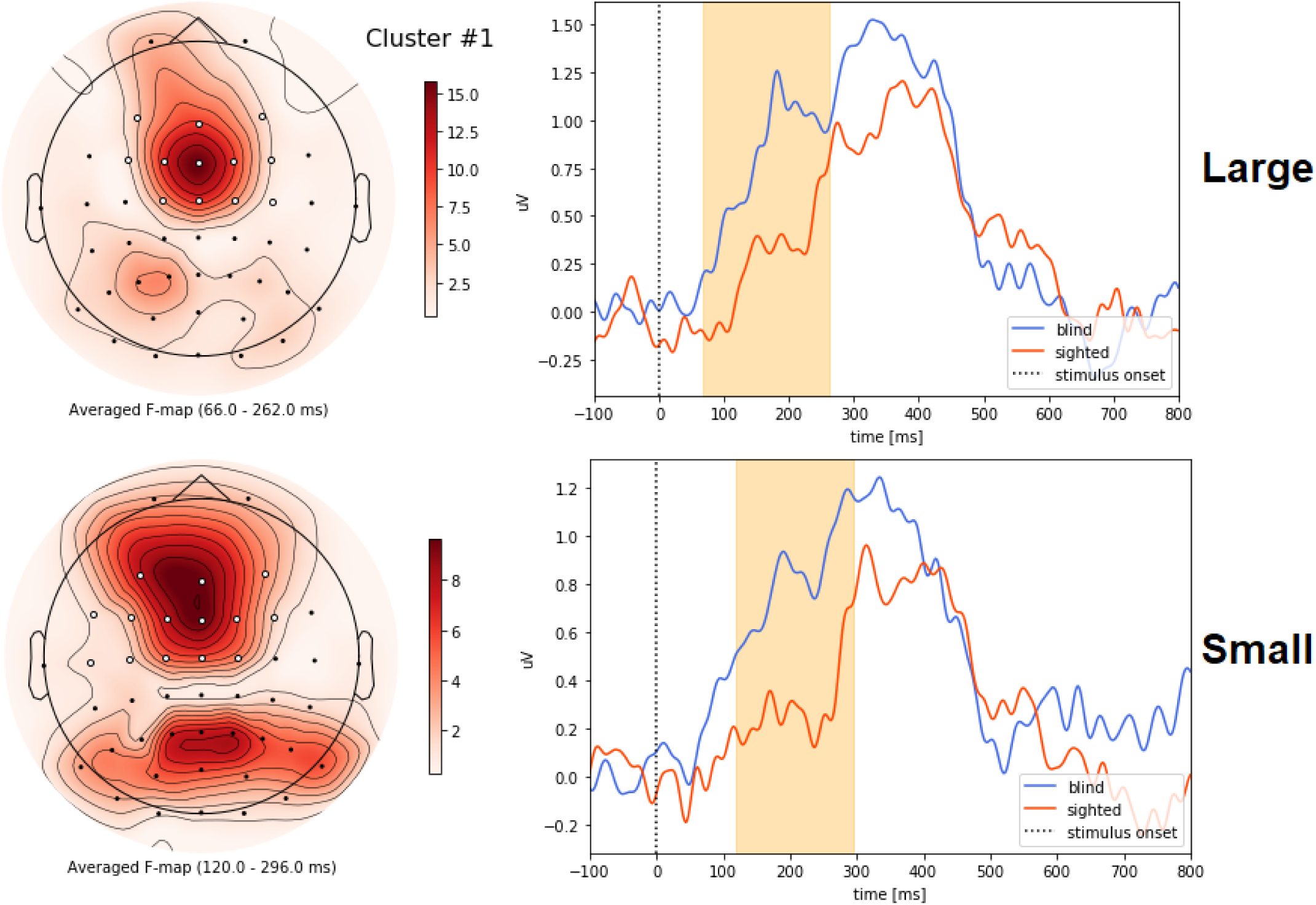
Significant differences from the cluster-based permutation test. On the right is the F-score map with highlighted significant channels. On the left is the average waveform for the cluster. The colored area on the waveform plot corresponds to the *p* < 0.05 significance window.

### Classification

For BCI with 8 stimuli, the chance accuracy level is 12.5%. In the experiment, both large and small stimuli conditions exceeded this threshold for all records. Median within-subject multiclass accuracy with 10 simulated target stimuli repeats for large stimuli was 0.75, and 0.53 for small stimuli. The tendency of smaller stimuli yielding less is still present when the sample is divided into the blind and the sighted groups: in the sighted group reconstructed accuracies for large and small stimuli were 0.62 and 0.5, and in the blind group 0.95 and 0.77 correspondingly. This result was statistically significant for the combined sample (Wilcoxon 𝒲 = 27, *p* = 0.0036), for the blind group (𝒲= 8, *p* = 0.047), and for the sighted group (𝒲 = 6, *p* = 0.03). It is worth noting, that despite lower classification accuracies for the sighted subjects and for smaller stimuli, one of the sighted participants was still able to achieve 100% target selection accuracy in both conditions.

The overall performance of the blind subjects was significantly better than that of the sighted ones with large stimuli(𝒰 = 15, *p* = 0.008) and in the combined sample (𝒰 = 115, *p* = 0.01). For small stimuli, the difference between the blind and the sighted groups was not significant (𝒰 = 37, *p* = 0.17), see Fig 6. Fig 7 represents the projected multiclass accuracy for the different number of stimuli presentations. The relation of the classification accuracies between groups and conditions remains the same for any number of stimuli presentations.

**Fig 6.**
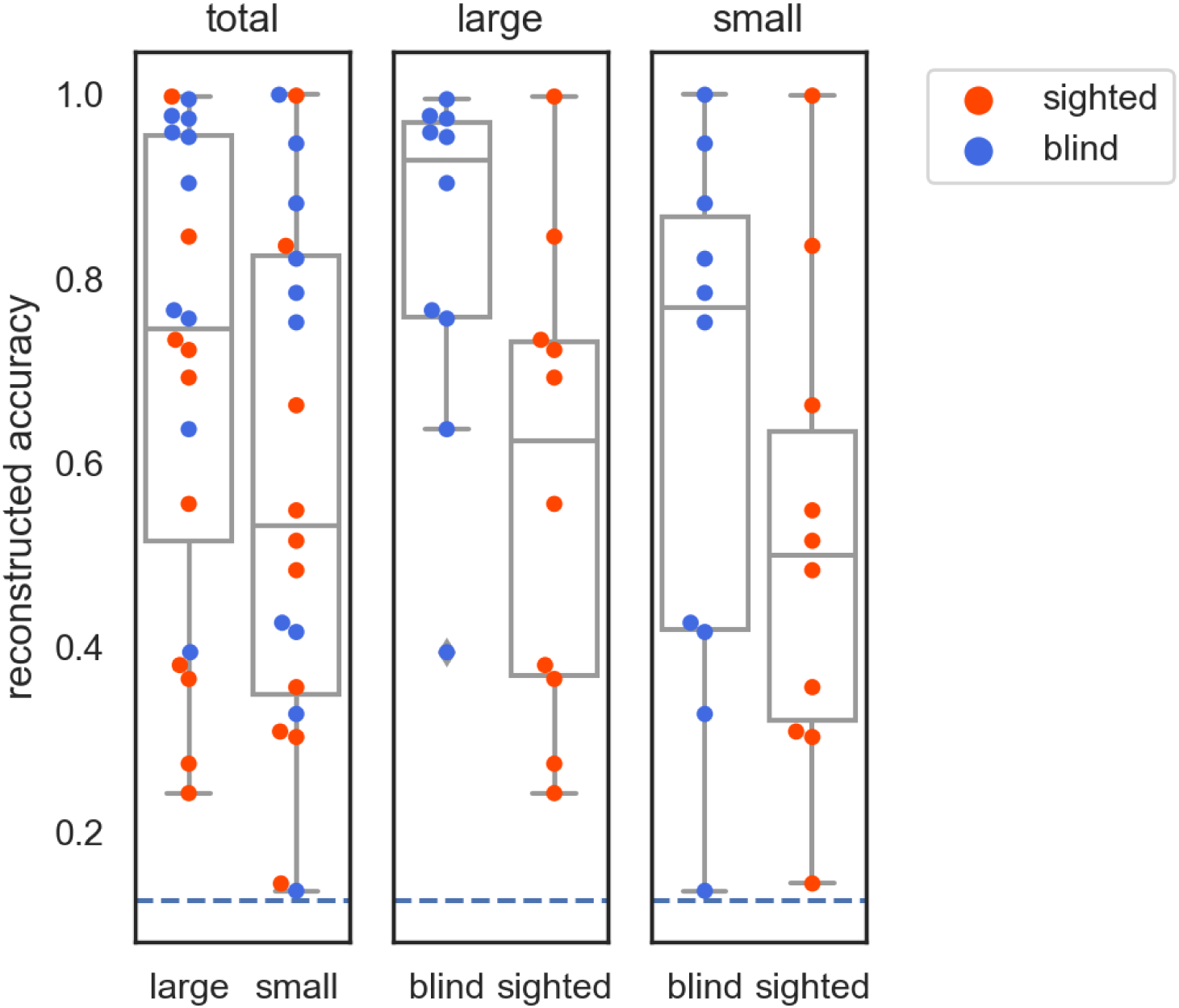
Reconstructed classification accuracy for all stimuli conditions. Median, 25%-75% quartiles, range, data points. The dashed line represents the chance level.

**Fig 7.**
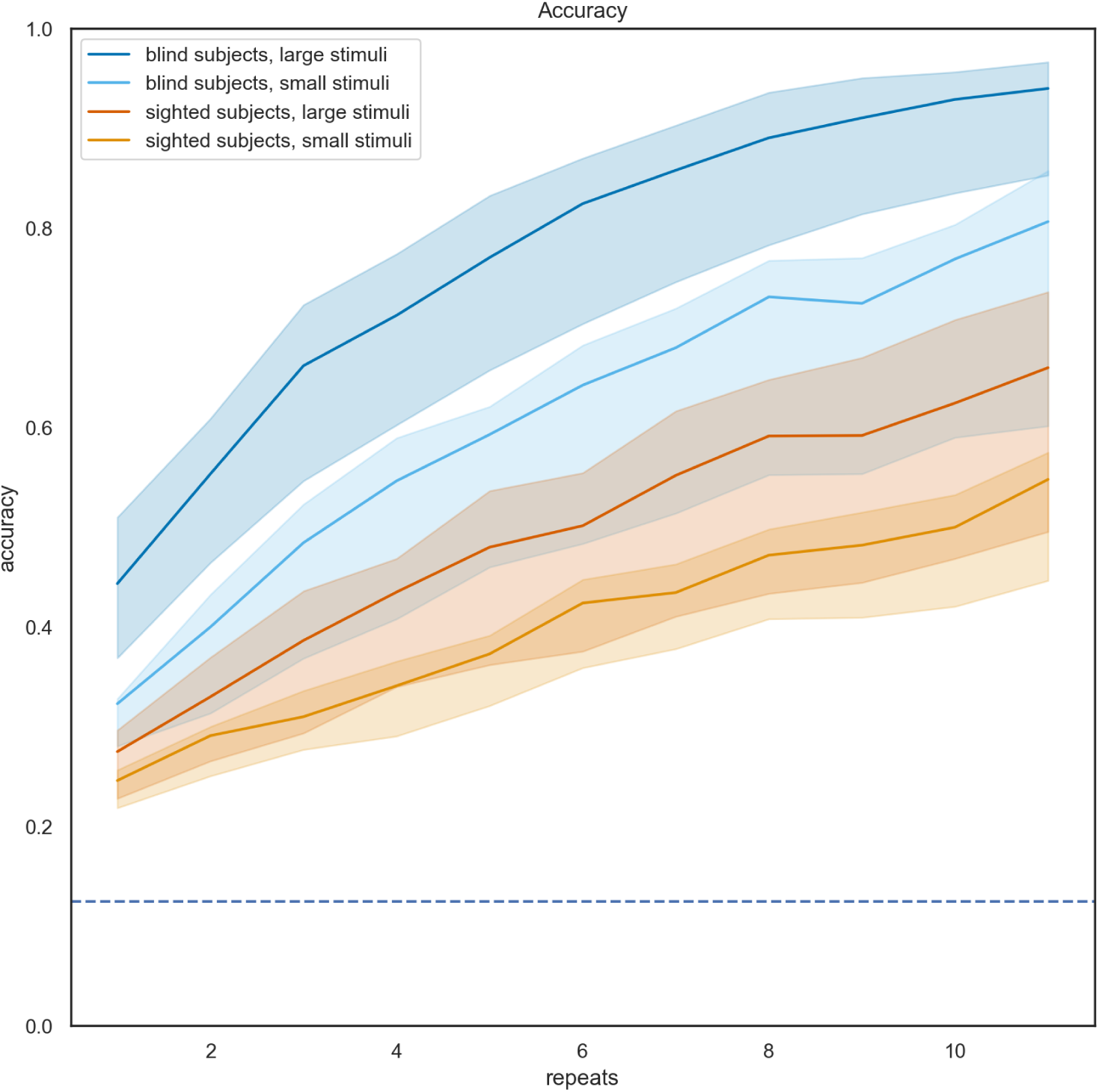
Reconstructed classification accuracy for all stimuli conditions and subject groups. Lines represent medians, areas represent 25%-75% quartiles. The dashed line represents the chance level.

The classification accuracies in two stimuli conditions are highly correlated within subjects both in the combined group of the blind and the sighted subjects (*τ* = 0.64, *p* = 3 × 10^−5^) and in each subgroup: the blind (*τ* = 0.5, *p* = 0.047) and the sighted (*τ* = 0.77, *p* = 0.001), see Fig 8. No significant correlation was found between subject age and classification performance.

**Fig 8.**
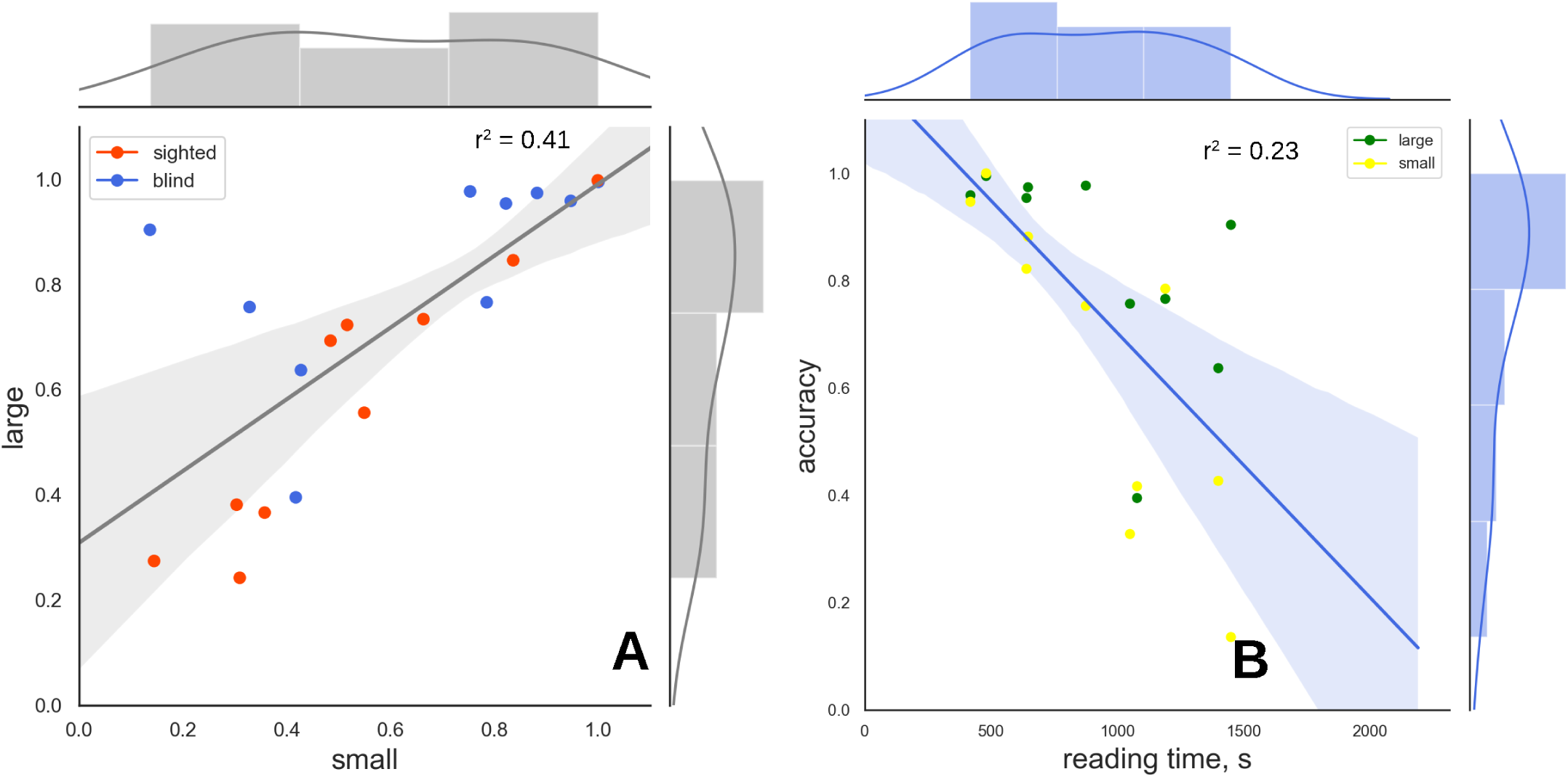
Correlations of classification accuracy. (A) Correlation between each subject’s performance in all conditions. (B) Correlation between reading speed and classification accuracy for the blind subjects for all stimuli types.

The accuracy in the blind group is significantly correlated with Braille text reading time, measured at the beginning of the experiment (*τ* = −0.47, *p* = 0.004). Interestingly, there is no significant correlation between reading speed and self-reported daily Braille time, and the latter is not correlated with ERP classification accuracy either. Using the same classification pipeline, we were able to classify the blind from the sighted by single-trial ERPs with 62% accuracy, which is another metric of differences in waveforms, that is reported earlier.

## Discussion

The cluster-based permutation test does not allow for highly specific claims about the significance of the effect in space and time [30]. However, the approximate frontocentral location of the cluster and its timing at around 100-300 ms poststimulus do fit into the early cognitive processing stages of the stimuli. This result partially reveals some possible mechanisms of sensory compensation, at least in the perceptual task that we have studied. Other possible outcomes would have included differences located in post-P300 late cognitive processing or differences in non-target evoked potentials.

It has been established in the seminal paper by Sadato [35], that visual cortex is activated in Braille readers, with Cohen [36] showing that this activation not just occurs, but also is necessary for haptic object recognition. We were not able to find any electrophysiological markers in the occipital areas, differing between the blind and the sighted. This may be due to the specificity of the BCI task, requiring discrimination of different fingers. Previous studies have shown that visual areas are specifically activated during tasks, requiring recognition of Braille letters, but not by nonsense Braille combinations or touching reading hand [37]. Recently, it has been shown that the sighted subjects can experience similar cortical reorganization after mastering Braille reading [38]. There is evidence, that both in the blind and the sighted subjects visual cortex can be recruited to recognize some tactile stimuli, at least Braille letters.

Perhaps, more complex BCI paradigms, involving discrimination between Braille letters, would be capable of utilizing visual areas, possibly enhancing BCI performance. However, while the classification of simple easily differentiable Braille-like shapes which has been shown in sighted subjects [7] the discrimination of complex shapes, presented on Braille display, can be hard even for the blind subjects, since the perceptual task is very different from the sliding finger movement that is usually used in reading scenarios. Successful classification may require longer stimuli presentation time.

Further research in this area may be directed in investigating the blind subjects in rapid serial tactile presentation BCI, requiring discrimination of different shapes under a single finger. Interestingly, no differences in classification accuracies between different fingers were found, despite the majority of participants being single-finger readers. Even multi-finger Braille readers are reported to almost never use several fingers to perceive different letters at the same time - when one reading finger is positioned on the letter, another is usually positioned on the space between words.

Lower performance in the condition with small stimuli can be partially addressed to subjects temporarily losing target cell, as well as the characteristics of stimuli themselves. Further evaluation of the effect on the size of stimuli may require a device with the fixed relative position of finger and Braille cell. Another option is to analyze kinematic data, tracking fingers during natural reading, a paradigm being used to study language processing in the blind. Participants did frequently report, that some Braille letters are easier to discriminate between than others, with reasons involving the size of the letter.

Cross-validation testing of the classifier on the recorded data allows not only to determine the usability of BCI but also to assess the significance of differences between samples with minimal *a priori* information. BCI performance in the within-subject learning paradigm depends mainly on the dispersion of event-related potentials within a single target selection session and the stability of differences between target and nontarget classes. During online classification, the performance also depends on the distribution similarity between learning and target selection samples.

ERPs depend on the level of fatigue, overall perceptual complexity of the task for the user, and cognitive workload [39]. However, it is still not obvious how changes in EEG translate into changes in behavior. This is why the higher BCI performance and even higher ERP amplitude in the blind group by itself are not enough to claim higher tactile acuity without additional behavioral tests. The correlation between Braille reading speed and the BCI performance (hence, ERP waveform characteristics) is an important result, possibly establishing a link between BCI classification accuracy and actual tactile acuity in a familiar task, at least in the blind group.

In this study, we have not found any correlations of BCI performance with age, while it has been reported, that older subjects demonstrate inferior performance in tactile BCI [40], and ERP BCI in general. Also, no correlation was found between subject age and Braille reading speed.

Previous research has shown that even in the same group of participants the blind subjects may be able to outperform the sighted subjects in one tactile test, but at the same time not differ in others [19]. Additional research would be needed to establish the mechanisms under the differences in tactile perception not only between the blind and the sighted but also inside the blind group.

Interestingly, in the present study, the success in the tactile task of Braille reading was correlated with success in completely different tasks of finger discrimination in BCI, despite what was claimed before [23]. It’s not obvious, how the ability to discriminate between spatial patterns with one finger can translate to better discrimination of similar stimuli under different fingers. There are possible mechanisms of irradiation of enhanced tactile acuity with training. For example, there have been reports of the transfer of spatial discrimination enhancements from hand to face [41] with possible mechanisms including Hebbian passive learning, but the fact itself asks for more replication and its neuronal mechanism needs to be clarified. Importantly, most previous studies that use metrics of Braille proficiency, did use self-reported metrics. We have demonstrated, that, at least in our cohort, the self-reported metric does not correlate with actual Braille reading speed. This may encourage the usage of objective metrics, like reading speed, especially since it can be integrated seamlessly into the experiment by Braille printing official documents regarding the study.

Differences in BCI performance that had been shown in this study once again brings the discussion about the importance of subject selection in BCI research. Here, we have significant differences between the target demographic (blind subjects) and the group that is mostly used for BCI research (healthy sighted subjects). Even more, some within-group dispersion in the blind subgroup is correlated with the ability to read Braille text. It is appropriate to ask, whether differences in BCI performance in other experiments (or even the phenomenon of BCI illiteracy) can be traced back to specific measurable subject characteristics that are not assessed during typical BCI study. While there is no comparable published data for visual BCI, some groups with specific sensory experience or training may be able to achieve different BCI performance from others. This opens possibilities for research in basic human neuroscience, but also should be taken into account when collecting public data sets or designing BCIs for practical use.

Further investigation is needed, with more focus on the connection between Braille literacy and performance in different haptic tests. The cause and effect relationship between tactile acuity and Braille proficiency is also an important question, having important scientific and social implications. With the advance and dissemination of speech synthesis, more blind people can process information without the need to use Braille. Combined with the high costs and difficulties in teaching Braille, this has led to decrease in Braille literacy, which some refer to as “Braille crisis” [42–44]. It is important to understand, to what extent the higher tactile acuity of the blind may be caused by extensive Braille experience, and to what extent it contributes to the quality of life.

## Conclusion

This is the first study to evaluate tactile BCI performance in the blind population. We demonstrated differences in evoked responses between the blind and the sighted cohorts and higher classification accuracy in the blind subjects. Future studies may improve BCI performance by fine-tuning to target demographics. A positive correlation between BCI performance and Braille proficiency opens a wide field of study regarding the causation of the effect.

## Supporting information

### Data availability

Anonymized raw EEG dataset in BIDS format [45] is available at https://gin.g-node.org/eegdude/BrailleBCI

### Code availability

Complete data analysis and processing pipeline is available at https://github.com/eegdude/erp_analysis/tree/brlbci

## Acknowledgments

Authors thank the “Con-nection” Deaf-Blind Support Foundation for the help with recruitment of blind subjects. Authors also thank Alexander Toroptsev and Svetlana Kim for the help with conducting experiments, Anna Vasilevskaya for valuable comments and Daniil Kuznetsov for language editing.

